# Identification of residue inversions in large phylogenies of duplicated proteins

**DOI:** 10.1101/2021.11.04.467263

**Authors:** Stefano Pascarelli, Paola Laurino

## Abstract

Connecting protein sequence to function is becoming increasingly relevant since high-throughput sequencing studies accumulate large amounts of genomic data. In order to go beyond the existing database annotation, it is fundamental to understand the mechanisms underlying functional inheritance and divergence. If the homology relationship between proteins is known, can we determine whether the function diverged?

In this work, we analyze different possibilities of protein sequence evolution after gene duplication and identify “residue inversions”, i.e., sites where the relationship between the ancestry and the functional signal is decoupled. Residues in these sites are masked from being recognized by other prediction tools. Still, they play a role in functional divergence and could indicate a shift in protein function. We develop a method to specifically recognize residue inversions in a phylogeny and test it on real and simulated datasets. In a dataset built from the Epidermal Growth Factor Receptor (EGFR) sequences found in 88 fish species, we identify 19 positions that went through inversion after gene duplication, mostly located at the ligand-binding extracellular domain.

Our work uncovers a rare event of protein divergence that has direct implications in protein functional annotation and sequence evolution as a whole. The developed method is optimized to work with large protein datasets and can be readily included in a targeted protein analysis pipeline.

## Background

Proteins perform their function either through protein-protein interactions, protein-ligand interactions, or catalyzing chemical reactions. At the molecular level, residues of a protein interact with a counterpart, namely a ligand (small molecule, protein, DNA/RNA, etc.). Herein we define such interacting residues as functional residues. The importance of predicting functional residues in a protein is evident as these residues can contribute to designing new functions, switching specificities, defining protein families and subfamilies, or identifying the occurrence of a functional innovation (e.g., a change of ligand specificity). Crystal structures in which the protein of interest was co-crystallized with its ligand, can readily identify functional residues. However, when the structure is not available, the identification of functional residues is not trivial. Previous attempts used the evolutionary information found in protein sequences and their homologs (1-3), approaches now facilitated by the global-scale genome sequencing effort driven by the development of high-throughput sequencing technologies. The prediction of functional residues by such methods is hampered by the presence of neutral mutations, namely amino acid substitutions that are neither beneficial nor disadvantageous (4). Non-synonymous neutral mutations are on average 10 times more abundant than advantageous mutations (5). While neutral mutations help to determine the phylogenetic position of a protein, mutations of functional residues are a signal of functional shifts that might occur independently of the phylogeny. When the relationship between phylogeny and function is decoupled, neutral mutations may mislead homology-based prediction methods, which are the most common way of functional prediction (6).

An event that often decouples the phylogenetic and functional signal is gene duplication. After gene duplication, two proteins follow a semi-independent evolution. For example, before diverging, the two duplicates may influence each other by gene conversion (7) or homomeric-heteromeric interactions (8), and they tend to diversify in the expression profile (9, 10). Gene duplication is prevalent in all domains of life (11), and often the duplicated proteins are reported to go through functional diversification (12). For example, in an event termed “sub-functionalization”, a protein with multiple functions (e.g., cellular receptors binding multiple ligands) might split its functions between the two gene copies after duplication. Previously, McClintock *et al*. showed that, in zebrafish HOX genes, the subset of functions inherited by the duplicated copies is different between fish and mouse – a phenomenon named “function shuffling” (13). In cases alike, the phylogenetic signal is misleading when used to predict the function of a “shuffled” orthologous protein. However, if the functional divergence is correctly identified, it allows to highlight the functional residues responsible for this transition, with reduced noise from the neutral variants. In this work, we address the identification of protein functional residues that are mutated during this type of functional rearrangement.

Residues responsible for a change of function within a protein family are usually called Specificity Determining Sites (SDS). SDS can be predicted by multiple methods (14). SDS prediction methods use the Multiple Sequence Alignment (MSA) or the 3D structure of the protein of interest to calculate a score based on conservation (15-23), evolutionary rate (24-26), or 3D structure properties (27). Most of these approaches require the user to provide the correct groupings of the homologous proteins. When this information is missing, the groupings are made according to the ortholog classification obtained by manual or automatic partitioning methods (28-31). However, the SDS predictions in automatically partitioned orthologs showed a lower sensitivity (32), demonstrating that an incorrect grouping negatively influences the prediction. The grouping usually follows the ortholog conjecture, namely that orthologs are more conserved than paralogs (33, 34). When this is not true, the SDS prediction is hampered. Therefore, the power to predict functional residues is limited by our ability to track protein function on the phylogenetic tree when it is not linearly inherited by orthologs. In our work we address this problem by identifying a signal of functional transition that might prove to be useful when annotating orthologs.

EGFR (Epidermal Growth Factor Receptor) is a tyrosine-kinase receptor that activates multiple signaling pathways after binding one of the seven EGFR ligands (35). EGFR is broadly expressed (36) and plays a crucial role in several aspects of organismal development and homeostasis like cellular growth, differentiation, metabolism, and motility (37). In fish, two copies of EGFR were kept after the Teleost-Specific Genome Duplication (TSGD) event that occurred about 350 mya in the actinopterygian lineage (38, 39). Lorin *et al*. showed that both copies of EGFR might have been retained because they are involved in the complex process of skin pigmentation (40), which is under high evolutionary pressure in most fish. Furthermore, the extracellular domain of fish EGFR, responsible for binding multiple ligands, likely went through sub-functionalization (41). For these reasons, EGFR constitutes a perfect model to study uneven functional inheritance events.

In this work, we observe a scenario where the function of a protein is not linearly inherited across orthologs, and we identify the functional residues responsible for the shift of functions. Our goal is to develop an algorithm that highlights the signature of a putative inversion of function, as could be an inversion of residues between the paralogs within the same species. First, we obtain a simple theoretical model that describes the likelihood of a residue inversion in comparison to other outcomes. Then, based on the model, we develop an algorithm that identifies residue inversions in a phylogeny, and we apply it in the context of fish EGFR duplication. Finally, we validate the results using statistical scores and simulated evolution. Our analysis shows a new way to investigate an important and understudied outcome of gene duplication.

## Results and Discussions

### Theoretical model for residue inversion identification

First, we constructed a simplified model to describe the evolution of a residue through a protein phylogeny after gene duplication and speciation. In the model, a tree branch (b1–b6) could be either mutation (1) or no-mutation (0) state, while a leaf node (Xa, Ya, Xb, Yb) has the possible states 0, 1, or 2 depending on the number of mutations in the preceding branches (Figure 1A). A configuration of six branch states univocally leads to a configuration of four leaf nodes. The model uses two branch length parameters, pre-speciation (t1) and post-speciation (t2), to calculate the probability of each of the 64 branch configurations. Using the model and a set of matching rules between the leaves (Figure 1B), we assessed the probability of seven categories/scenarios of configurations (Figure 1C): *Conserved*, all four residues match; *Type 1* divergence, residues match in only one paralog group; *Type 2* divergence, residues match per paralog group but not per species; *Recent divergence*, all residues match except for one leaf node; *Inversion*, residues of opposite paralog group match but not per species; *Species-specific adaptation*, residues match per species, but not per paralog group; and *Non-conserved*, collecting all events that do not fall in any of the other categories. We used the following formula to calculate the probability of each category:

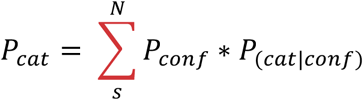

Where N are the 64 configurations of branch states, the probability *P*_*conf*_ of the branch configuration is given by the model (Figure S4), and the conditional probability for the category *P*_(*cat* | *conf*)_ is determined by the category’s matching rules.

**Figure 1.**
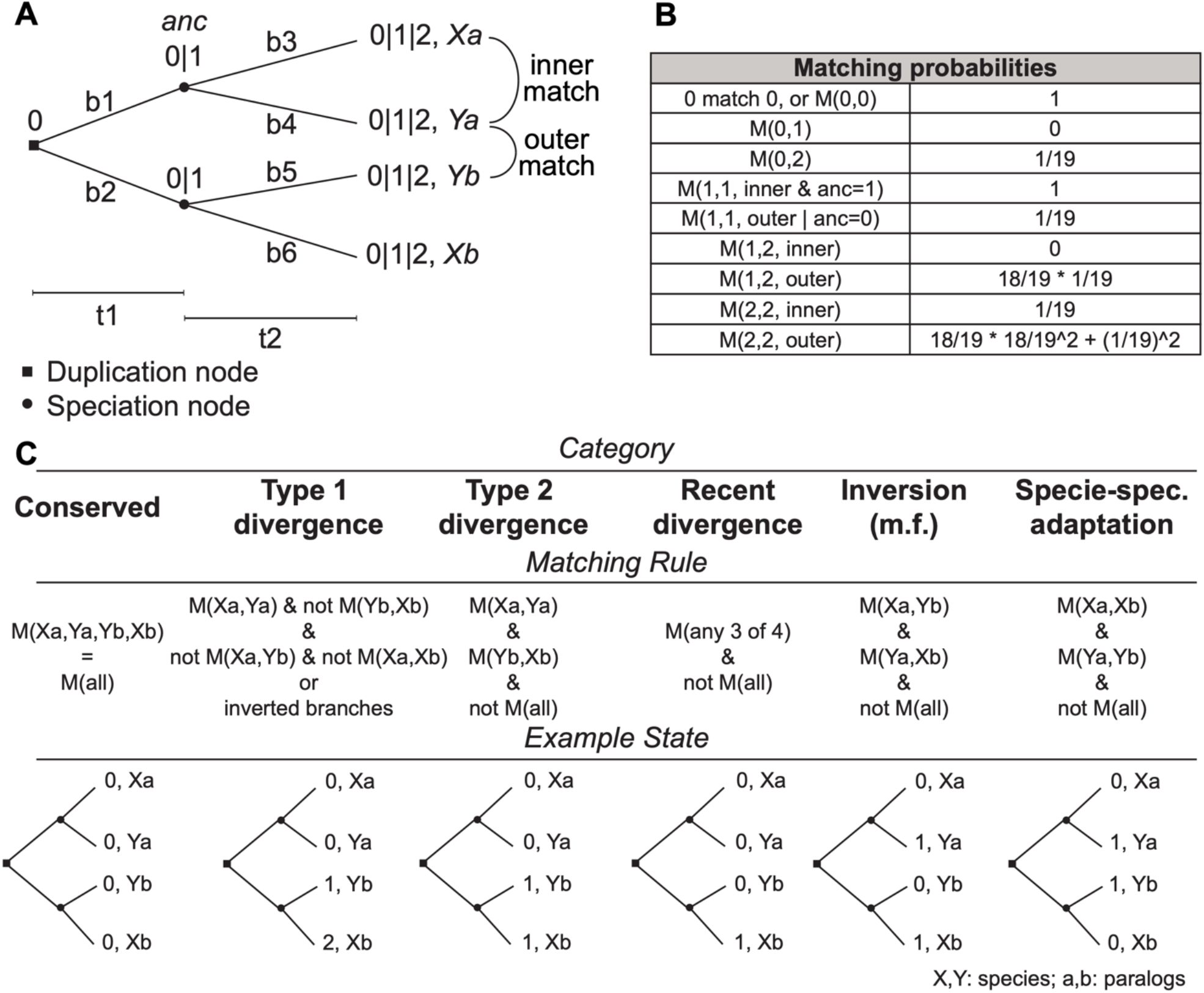
Theoretical model of the evolution of protein residue after gene duplication. **(A)** The structure of the phylogenetic tree that the model is based on. The branch lengths t1 and t2 are used to determine the probability of a mutation on each branch b1 to b6. A leaf node can be found in states 0, 1, or 2 depending on the number of mutations in the preceding branches. An inner match is defined to be a match between orthologs (Xa to Ya, or Xb to Yb), while an outer match is any other match. The probability for a match between two states is given by the table in **(B)** and represents the underlying transition to any of the 20 amino acids. **(C)** Description of the categories. The categories represent a biologically interpretable situation, suggested by their name. Given a certain outcome configuration of states, it is possible to calculate the probability of observing a certain category by using the matching rule. The “Example State” section shows the leaf configuration that gives the highest probability of observing the category described.

Next, we tested the probability of each category at varying branch lengths (Figure 2A). As expected, short branch lengths lead to a high probability of conserved residues, and long branch lengths lead to non-conserved residues. When the pre-speciation branch length is the longest of the two, we observe a high probability of Type 1 and Type 2 divergence. Whereas, when the post-speciation branch length is the longest, we observe a high probability of recent divergence. Interestingly, for any branch length combination, the two categories of inversion and species-specific adaptation have less than a 1% chance of appearing. From this analysis, we expect to rarely find these two events in real data.

**Figure 2.**
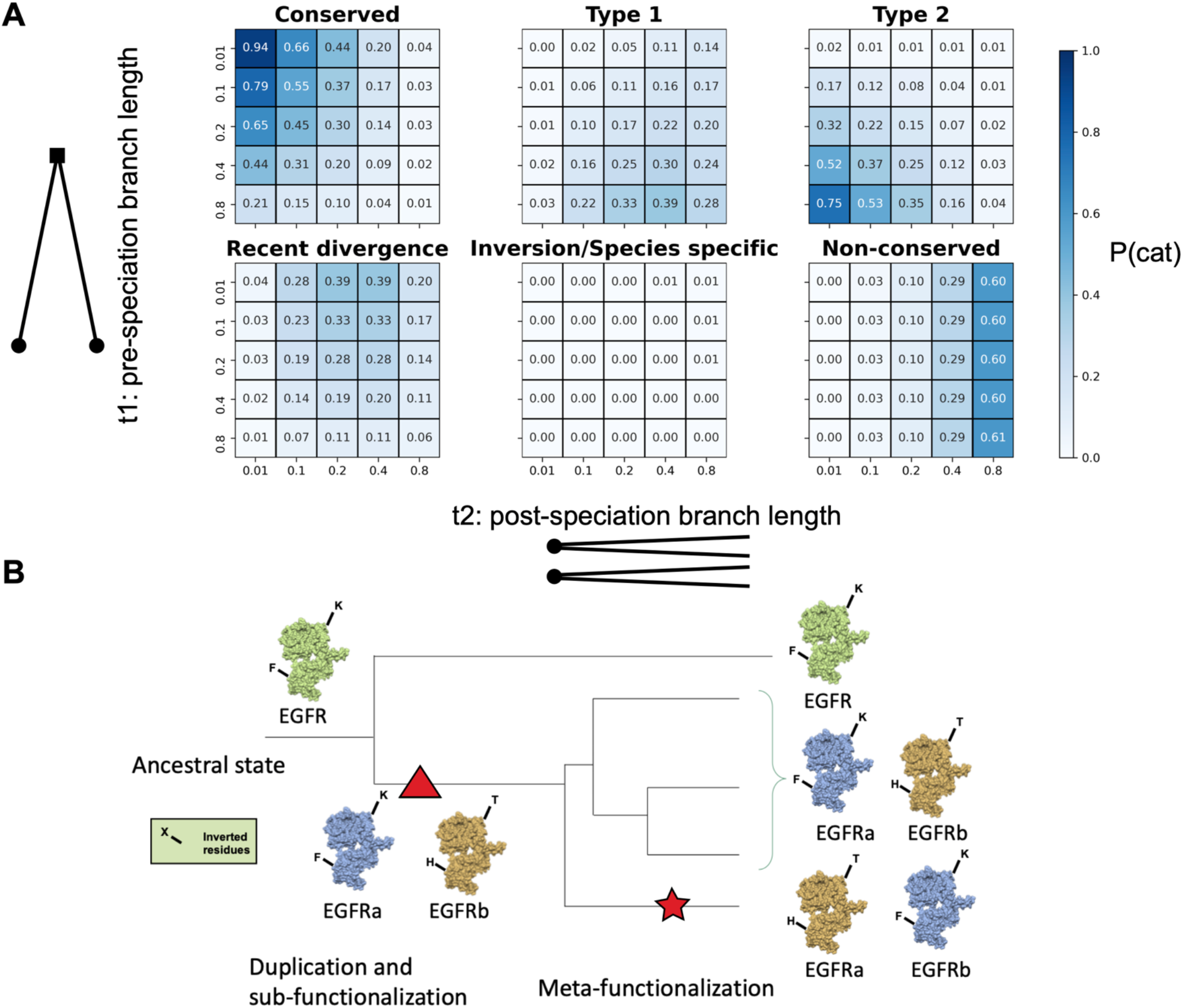
Theoretical model results. **(A)** The heatmaps show the category probabilities at different tree branch lengths as calculated using the theoretical model definitions. **(B)** Exemplification of “Meta-functionalization” (star), the putative driver of residue inversions in the phylogeny. A multifunctional protein (green) subdivides its functions (blue and yellow) between the two copies obtained after gene duplication (triangle). In a sub-group of species, the functional inheritance of the two copies is inverted. This event is revealed by the pattern of inverted residues compared to the majority of the other species.

From the previous results, residues inversions should not be recurrent in a phylogeny. Thus, an unlikely high presence of inverted residues may show that: 1) these residues are following a selective pressure directly related to the paralog function that, for a particular subgroup of species, is opposite and complementary to the other species; or 2) these residues act as compensatory mutations. A recurrent residue inversion could be the proxy of a functional rearrangement between the paralogs within a clade. A rearrangement of functions is facilitated in, but not limited to the Innovation Amplification Divergence (IAD) model of duplicates divergence (42), where paralogs partially retain their secondary functions. We describe this subgroup-specific functional inversion event as a possible outcome after the sub-functionalization of a duplicated gene and we suggest the name “meta-functionalization” from the Greek word “metathesis”, namely “put in a different order” (Figure 2B).

### Algorithm for the identification of residue inversions

We developed an algorithm to identify the events of inversion in a protein phylogeny using a multiple sequence alignment (MSA) and a phylogenetic tree. The algorithm was implemented as a python package named DIRphy, for the Detection of Inverted Residues in a phylogeny. DIRphy splits the protein sequences of the MSA into four groups according to the organism and orthology annotations, which can be either provided by the user or automatically done using a tree distance parameter. Based on the matching probabilities of the previous theoretical model, DIRphy calculates a score for each event of “Inversion” and “Species-specific adaptation”, representing its probability to occur (see methods for details). However, for this paper, we will focus only on the inversion events. The output of DIRphy is a list of positions above a defined threshold. When the organism grouping is manually selected, the script calculates both the observed and the expected probability of inversion between the specified groups in the given tree. Otherwise, when the organism grouping is not specified, the output table shows the observed probability of residue inversion given by the grouping that has the highest probability in that position. In the current version of DIRphy, only a binary paralog classification is allowed. DIRphy is released as an open-source project in Github: https://github.com/OISTpasca/protein-inversions

### Construction of a fish EGFR dataset and identification of residue inversions

We tested DIRphy in the phylogenetic tree of the Epidermal Growth Factor Receptor (EGFR) in the fish lineage. First, we filtered 88 fish genomes (taxon 41665) from the European Nucleotide Archive (ENA) (43) to obtain a dataset of 167 fish EGFR protein sequences. The dataset included all high-quality (N50 > 1Mb) teleost genomes plus one outgroup before the TSGD (spotted gar). From the phylogenetic analysis of this dataset (Supplementary figure 1), three clear duplication events can be observed. The first, most ancient duplication coincided with the TSGD, around the time of the split with gars (∼350 mya), and resulted in the separation of the two copies: EGFRa and EGFRb. Two more copies were found in most salmonids, corresponding to the salmon-specific whole-genome duplication around ∼80 mya (44), and one more copy in goldfish, possibly due to the carp-specific whole genome duplication about ∼10 mya (45). The longer branch lengths of EGFRb (Mann-Whitney test, p-val: 2.305e-11) indicate that EGFRb is evolving more rapidly than its counterpart. Furthermore, the EGFRb gene was more commonly lost. Out of 15 gene loss events, only one species lost EGFRa (*S grahami*), while 14 species lost EGFRb.

Next, we tested the previously computed phylogeny and MSA of fish EGFR for residue inversions using DIRphy. We decided to compare the Cypriniformes clade with the other teleosts because of the high coverage of genomes in both groups, and a sufficiently long separation between the two groups to allow functional divergence on the protein sequence (Figure 3A). We observed a distribution of the scores that resembles an extreme-value distribution, with most of them below 0.01 (Figure 3B). Using the theoretical model, we calculated that the expected value for the probability of residue inversions in the same tree is 0.002, much lower than the score of the 99th percentile of 0.16. This percentile value was used as a threshold to select eight sites, the majority of which (six) were from the EGFR extracellular domain. We highlighted the positions of the six sites on the 3D structure of EGFRa and EGFRb extracellular domain from a representative Cypriniformes (*S. anshuiensis*), modeled using the human structure template (Figure 3C). Of the six selected positions, only one (MSA pos 520) was found at the ligand-binding pocket interface. Pos 520 corresponds to Phe-357 in hEGFR. Previous studies showed that the hydrophobic interaction between Phe-357 and Tyr-13 in the ligand hEGF is determinant for the binding (46). In the fish clade *Xiphophorus*, the observed change between Phe and His at this position is considered to be the determinant cause of the different responses of EGFRa and EGFRb after ligand stimulation (47). Out of the six positions in the extra-cellular domain, two showed a conservative substitution (Figure 3D). MSA position 506 contains hydrophobic and aliphatic amino acids (Ile or Val), while position 491 shows a small and uncharged amino acid (Ser or Thr). All other positions exhibited a shift of amino acid physicochemical properties. These results show that DIRphy can identify inverted residues in a protein duplication phylogeny, regardless of the amino acid substitution type.

**Figure 3.**
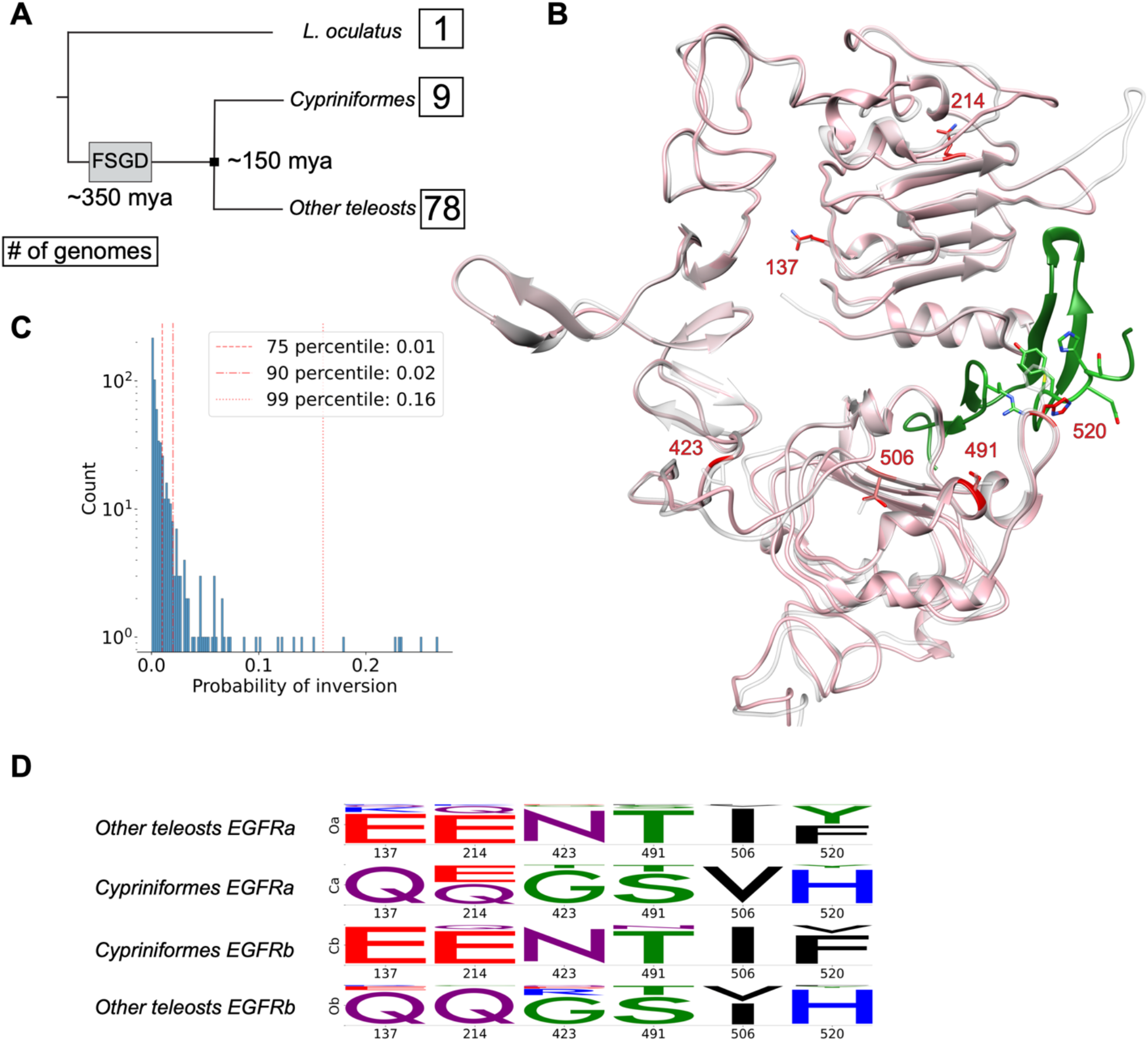
Inverted residues in fish EGFR. **(A)** Schematic representation of the fish dataset phylogeny. The dates (mya: million years ago) indicate the time of the Fish Specific whole Genome Duplication (FSGD) and the separation of Cypriniformes fish to all other teleost fish. The number in the boxes represents how many genomes are in the dataset for that group. **(B)** 3D model superposition of *S. anshuensis* EGFRa (pink) and EGFRb (white), generated by homology using human EGFR as a template (1IVO) (48). The inverted residues have been highlighted in red. The ligand EGF (green) was taken from the human model after superposing the receptors. **(C)** DIRphy score distribution. The residue inversion probability score was calculated for each site in the MSA that has less than 60% gaps. The top 1% of sites were further characterized. **(D)** Logo representation of the four sub alignments (two species groups, two protein copies) in the inverted residue sites. The logo represents the normalized amino acid count per column and was obtained using the python package Logomaker (49).

### Score validation by simulated evolution

We statistically validated the score observed in the fish EGFR data using a simulated evolution experiment. In this simulation, random starting amino acids are run through a phylogenetic tree that has the same topology as the previously computed fish EGFR tree. The simulation used the same evolutionary model of the fish EGFR tree to output a MSA as a result. Compared to the fish EGFR MSA, the simulated evolution MSA showed on average lower DIRphy scores while having a similar shape of the score distribution (Figure 4A). No specific amino acid was found to have high scores. Interestingly, the three residues involved in the interaction in position 520 (His, Tyr, and, except for one site, Phe) failed to produce any score higher than 0.05 in the simulation (Figure 4B). For further analysis, we used the 99^th^ percentile score of the simulation as a threshold for selecting inverted residues in the real dataset (Supplementary table 1, Supplementary figure 2). In summary, the simulated evolution experiment provided a score threshold for detecting residue inversions and confirmed the low chance of this event in the fish EGFR dataset, as observed in the theoretical model.

**Figure 4.**
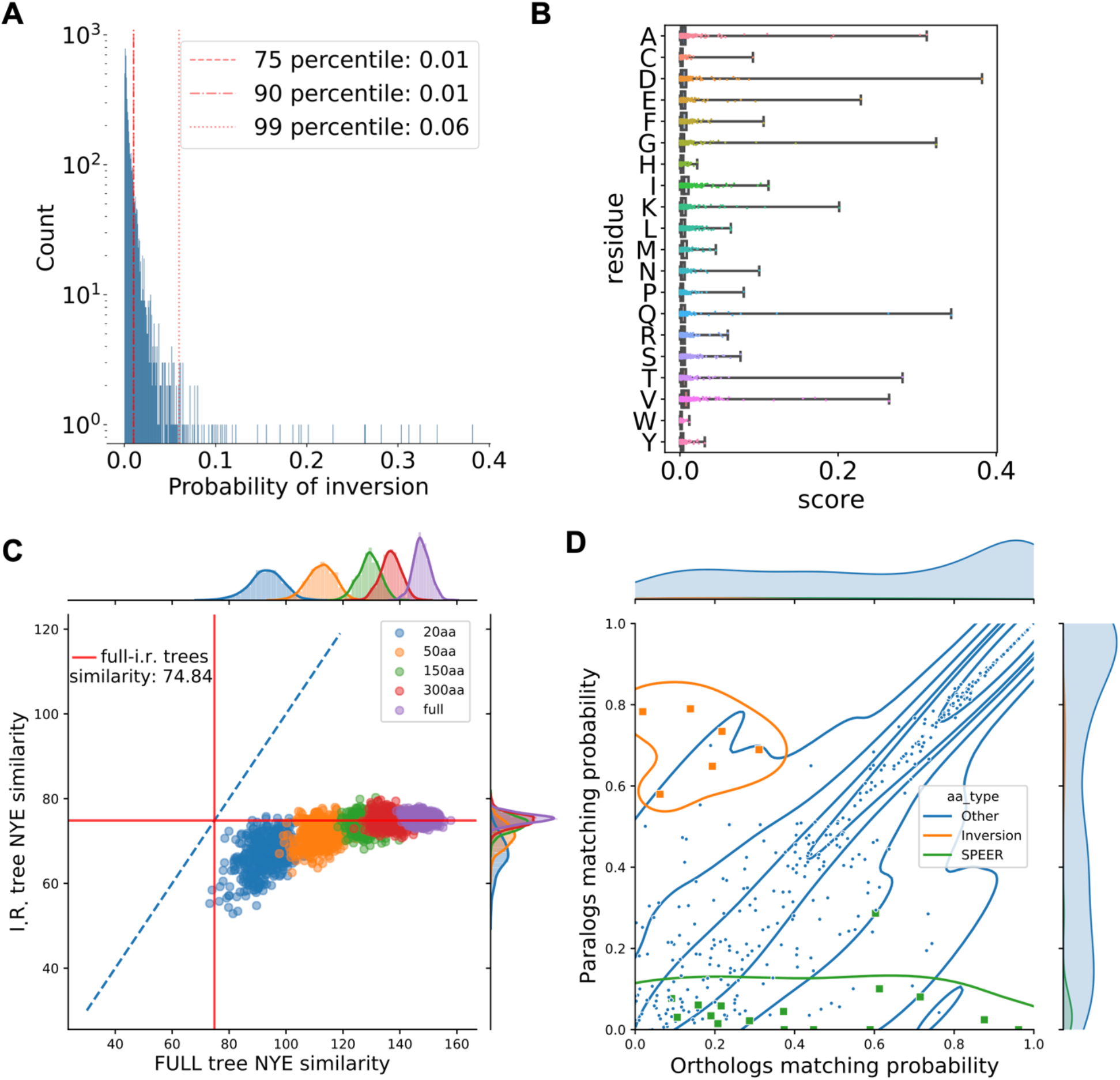
Outcome to the validations. **(A)** Distribution of the simulated evolution DIRphy scores. A 5000 random amino acid sequence was evolved through the fish EGFR phylogenetic tree using the same evolutionary models used to generate the tree. The resulting MSA was used to compute the DIRphy score. **(B)** Distribution per amino acid of the simulated evolution DIRphy scores as shown by the reference (*S. anshuiensis* EGFRa). **(C)** Bootstrap trees similarity to the full and inverted residue trees. The color represents the length of the sub-alignment used to generate the bootstrap tree. The red line shows the similarity between the full and inverted residue trees. The blue line is the identity line. **(D)** Comparison of the sites identified by DIRphy and SPEER. The matching probability of the HMM of four sub-alignments was used to compare between species the orthologs (EGFRa vs EGFRa, EGFRb vs EGFRb) and paralogs (EGFRa vs EGFRb). The matching probability is calculated as the average of two dot products of the frequency arrays. The orange color shows sites where a residue inversion was identified, while the green color shows sites where the p-value of SPEER score is lower than 0.01.

### Tree bootstrap

We characterized the phylogenetic information carried by the residue inversions when reconstructing the correct fish EGFR phylogeny tree structure. First, we computed a phylogenetic tree using the sub-alignment of 19 inverted residue sites that score higher than the 99^th^ percentile in the simulated evolution experiment (Supplementary figure 3). In this phylogenetic tree, we can still see two defined groups of sequences corresponding to the two copies of EGFR. Though, as expected, all the sequences from Cypriniformes EGFRa cluster together with the EGFRb group, and vice versa the Cypriniformes EGFRb cluster in the EGFRa group. When we compared this tree to the full-alignment tree, we calculated a similarity value of 74.84 out of 165 using the tree similarity score of Nye *et al*. (50). Next, we generated a pool of bootstrap trees with a reduced alignment length, and we checked their similarity to the full alignment and the inverted residue trees (Figure 4C). We observed a decrease in tree similarity to the full tree proportional to the decrease in alignment length. However, the inverted residue tree distance is statistically less similar to the full alignment tree than the bootstraps, even among the bootstraps with an equivalent length of 20 residues (student t-test p-value: 0.0019). This result suggests that the inverted residues are just a minority of the sites in the alignment and that they disperse the phylogenetic information faster than the average. This view is compatible with the hypothesis that a few functional mutations are hidden behind an overwhelming amount of neutral or nearly-neutral variants, complicating their detection.

### Comparison of DIRphy inverted residues to Specificity Determining Sites (SDS) prediction

We predicted the SDS of fish EGFR using SPEER (32), and observed a marked difference in the type of identified positions compared to DIRphy. To compare the two methods, we defined a “matching probability” as the dot product of the amino acid emission probability vectors of two HMM models. Then, we compared, for each site of the fish EGFR alignment, the mean matching probability between HMM models of the orthologs (cypriniformes EGFRa to other fish EGFRa, and cypriniformes EGFRb to other fish EGFRb) with the mean matching probability between the paralogs (cypriniformes EGFRa to other fish EGFRb, and cypriniformes EGFRb to other fish EGFRa) (Figure 4D). From this analysis, it is evident that SPEER tends to predict sites with a high ortholog-low paralog correlation. These sites are likely to possess a paralog-specific function. On the other hand, DIRphy identifies diametrically opposite sites, with a high paralog-low ortholog correlation. Both types of sites deviates from the diagonal, a pattern that suggests a functional adaptation. However, the sites that show an inversion are more challenging to identify for SDS predictions because, for a subset of species, the conservation pattern is inverted, and the signal is averaged out. In summary, the identification of residue inversion event has the potential to improve functional residue predictions, as DIRphy is able to identify functional sites that are overlooked by SDS prediction methods.

## Conclusions

In conclusion, we have detected an event in paralogs that lead to the inversion of functional residues. This new event has been described by a theoretical model and validated by literature and bioinformatics studies. To the best of our knowledge, we are the first to identify and describe the residue inversion event, possibly because this event might be rare. However, residue inversions are potentially exploited for functional divergence and, if missed, might lead to the wrong classification of proteins in the correct functional groups. We provide a general tool, named DIRphy, to identify residue inversions in a large protein dataset. DIRphy can be easily integrated in an existing pipeline of protein annotation to improve functional annotation and provide the positions that might have been overlooked by other functional site prediction methods.

## Materials and Methods

### Theoretical model

The model was built on the following assumptions: 1) equal branch length between the two paralogs: b1 = b2, b3 = b4 = b5 = b6; 2) only zero to one mutation can occur in each of the six branches; 3) after a mutation, each residue is equiprobable; 4) no selective pressure; 5) the probability of a mutation on a branch solely depends on the branch length (mutation rate) and is *P* = 1 −*e*^λ^ where P is the probability of a mutation and λ is the mutation rate.

Given the probability of a mutation in each of the six branches, we can calculate the probability of all the 64 (2^6^) configurations of mutations on the tree. A configuration unequivocally leads to a determined leaf node state (Supplementary figure 4). We defined the leaf node states with the allowed values zero, one, or two. The state value represents the number of mutations that happened in the branches connecting to the leaf node (Figure 1A). Then, we defined the probability of a match between leaf nodes based on the state value. The probabilities describe the situation in which a residue can mutate to one of 19 possible other residues. In some cases, the matching probability depends on the state of the ancestral node before speciation, e.g., a single or double mutation in the inner branch leaves. Finally, we defined seven categories based on the type of matching at the leaf nodes, and we defined their probability of occurrence based on the matching probabilities between leaf nodes.

### Model categories

Here we give a brief description of the model categories and formulas used to calculate their probabilities in the model.

#### Conserved

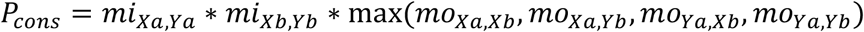

Where *mi* stands for the inner match, *mo* stands for the outer match and corresponds to the probabilities of a match in figure 1B. The conserved category collects the states where a site is invariant in all four leaf nodes. It could arise from no mutations, but also from (two, three, or) four mutations to the same amino acid. In the formula, the maximum value of the outer matches is used as the best approximation of the conditional probability of matching all leaf nodes given the two inner matches.

#### Type 1 Divergence

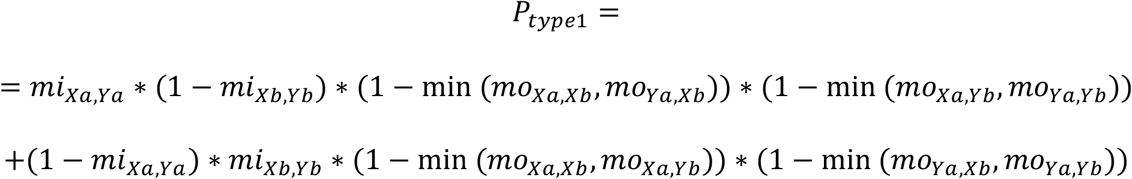

The type 1 divergence collects the states where the amino acid is matching between species only in one paralog, while there is no match in the other paralog.

#### Type 2 Divergence

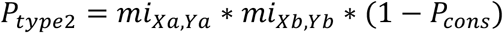

The type 2 divergence collects the states where the two paralogs display a different amino acid but are conserved between species. Type 1 and type 2 classifications are based on (51).

#### Recent Divergence

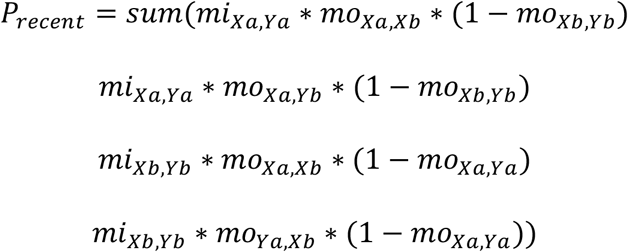

The recent divergence collects the states where only one leaf node is different (diverged) compared to the other three nodes.

#### Inversion / Specie-Specific Adaptation

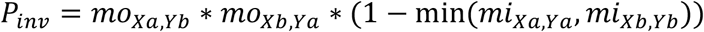

And similarly,

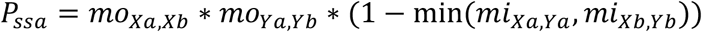

These two categories represent the states where the amino acid does not match between orthologs but matches between paralogs for the inversion or between species for the species-specific adaptation.

### Calculation of inverted residue score

We devised a score to identify inverted residues in a phylogeny. The score was based on the probability of observing an inversion in the previously described model and calculated with the following steps: 1) Divide an MSA into four sub-alignments (two EGFR copies and two species groups). 2) Generate four amino acid frequency arrays, optionally normalized by pseudo-counts*. 3) Calculate the probability of a match between two groups using the dot product of the frequency array. 4) Calculate the joint probability of inversion (or similarly for species-specific adaptation) from the conditional probabilities and the frequency array matching using the following formulas:

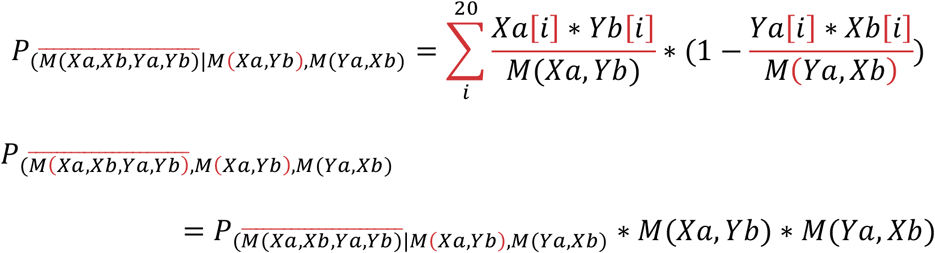

Where *Xa,Xb,Ya,Yb* are the amino acid frequency arrays for the MSA sequence groups with the same names, *i* is the counter spanning each amino acid. The probability of a match (e.g., *M(Xa,Yb)*) is given by the array dot product: 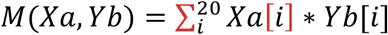 The latter joined probability represents the inverted residue score.

### * Pseudo-count normalization

To correct any possible bias given by groups with a small number of species, we implemented a pseudo-count normalization of the amino acid frequency array, as previously done in Tatsuov *et al*. (52). We used the LG protein substitution matrix (53) as background amino acid frequency probability. The value of the beta parameter for the pseudo-counts formula was set by default to five; however, it is possible to modify this parameter before running the pipeline.

### Fish dataset

To test the inverted residue score, we generated a fish genome dataset. First, we downloaded all genomes from the European Nucleotide Archive (ENA) belonging to taxon 41665 (Actinopterygii). Through this method, we obtained 88 fish genomes. Next, we downloaded the pre-annotated fish EGFR protein sequences from the ENSEMBL database (54) and used them to build an HMM profile with the HMMER package (55). The HMM profile was used as a query in Augustus package suite (56) to search for EGFR related genes in the fish genomes. We then filtered out interrupted CDS, sequences clustering with other ErbBs in a phylogenetic tree, and sequences with an aberrant branch length in the non-synonymous codon tree. After this procedure, we obtained 167 fish EGFR protein sequences.

### Phylogenetic analysis

We performed the phylogenetic analysis of the fish EGFR protein sequences using MAFFT (57) to align the sequences, and IQTREE (58) with ModelFinder (59) to search for the best evolutionary model to generate the phylogenetic tree. To generate the synonymous tree, we used paml CODEML (60).

### Sequence and structure analysis

The sequences and alignments were handled using Unipro Ugene (61). The protein structure images and analyses were performed with UCSF Chimera (62). The modelling of fish EGFR structures was performed using the SWISS-MODEL web server (63) and AlphaFold 2 (64).

### Simulated evolution

We ran a simulated evolution experiment using an in-house pipeline based on the Pyvolve python package (65). The pipeline simulated an evolutionary pathway of 5000 random amino acids on the fish EGFR phylogenetic tree, using the same model of evolution and evolutionary rates that were used to construct the tree (JTT with rate heterogeneity) (66-68). The simulation generated an output alignment that was used to run the DIRphy pipeline, to calculate the base probability of an inversion.

### Bootstrap

To perform the bootstrap, we used an in-house Matlab script. We selected from the fish EGFR DIRphy prediction the 19 sites with a score higher than the 99^th^ percentile of the simulated evolution scores. We calculated a phylogenetic tree for the full alignment and the inverted residues alignment using the neighbor-joining algorithm (69) and the BLOSUM80 matrix (70). The similarity distance between trees was calculated using the method described in Nye *et al*. (50). We then performed 500 bootstrap alignments for each set of alignment lengths: full, 250, 100, 20. We excluded columns with 90% or more gaps and repeated the sampling whenever one sequence did not have at least one non-gap position. For each bootstrap alignment, we generated a tree, then calculated the distance to the full tree and inverted residue tree.

## Declarations

### Ethics approval and consent to participate

Not applicable

### Consent for publication

Not applicable

### Availability of data and materials

The data and scripts used to support the conclusions of this article are available in the DIRphy Github python repository: https://zenodo.org/badge/latestdoi/346201641.

### Author contributions

S.P. and P.L. designed the project. S.P. performed the research. S.P. and P.L. analyzed the data and wrote the manuscript.

## Acknowledgments

We thank Federica di Palma, Tarang K. Mehta, and Wilfried Haerty for the thoughtful discussions on fish phylogeny. Stanisław Dunin-Horkawicz and Dan Kozome for critical reading of the manuscript. We are grateful for the help and support provided by the Scientific Computing and Data Analysis section of Research Support Division at OIST. Financial support by the Okinawa Institute of Science and Technology to P.L. is gratefully acknowledged.

