## Supplementary Data for "Identification of residue inversions in large phylogenies of duplicated proteins"

* Corresponding author

**Supplementary figure 1. Fish EGFR phylogeny.** Phylogenetic tree of the 167 EGFR proteins found in the fish genomes dataset. The label shows, in order, the name of the species where this EGFR was found, the contig name, start position, end position, the length in DNA bases, and the AUGUSTUS fastBlockSearch score, separated by underscores. The coloring shows how many genes are found in the species of this EGFR. The annotated sequences of zebrafish and tilapia EGFR were taken from the ENSEMBL database and added to the tree for reference. The main nodes bootstrap values are shown.

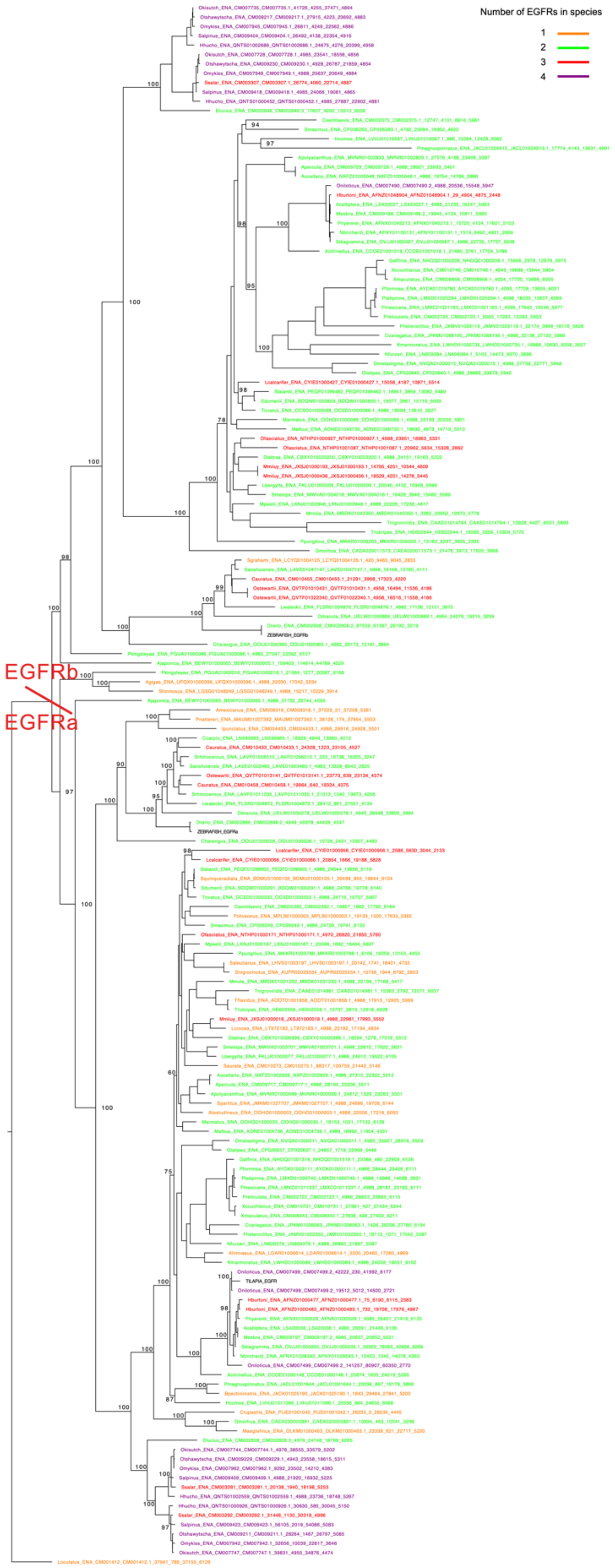

**Supplementary figure 2. Logo of the fish EGFR high DIRphy scoring inverted residues.**

The output of DIRphy for the sites of the phylogeny of fish EGFR in which the score is higher than the 99% percentile of the simulated evolution experiment (>0.07). Oa and Ob stands for Other fish EGFRa and EGFRb, while Ca and Cb stands for Cypriniformes EGFRa and EGFRb.

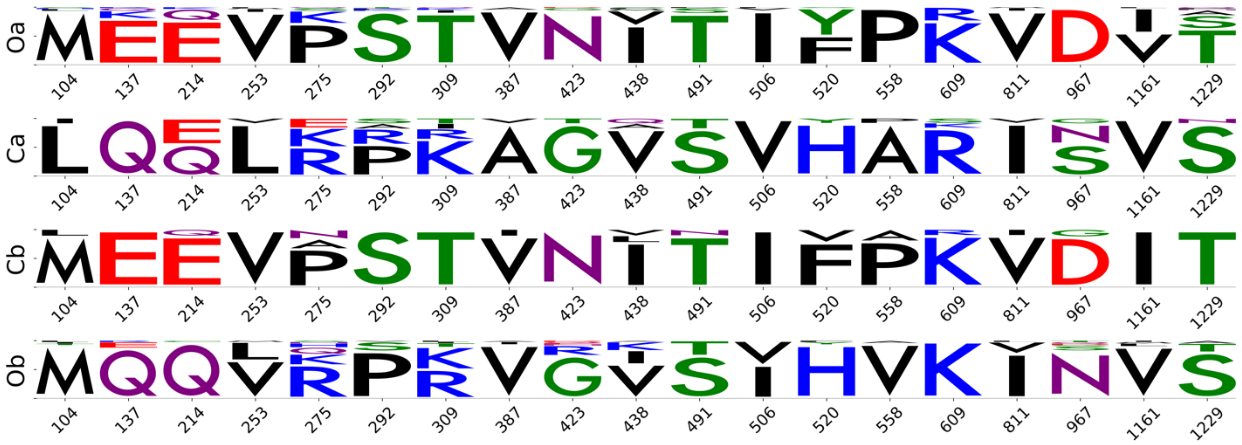

**Supplementary figure 3. Comparison of the inverted residues tree.** The two trees made from the full alignment (left) or the inverted residues sub-alignment (right) are colored by the four groupings used to calculate the DIRphy score: Cypriniformes EGFRa (red), other fish EGFRa (orange), Cypriniformes EGFRb (blue), other fish EGFRb (teal).

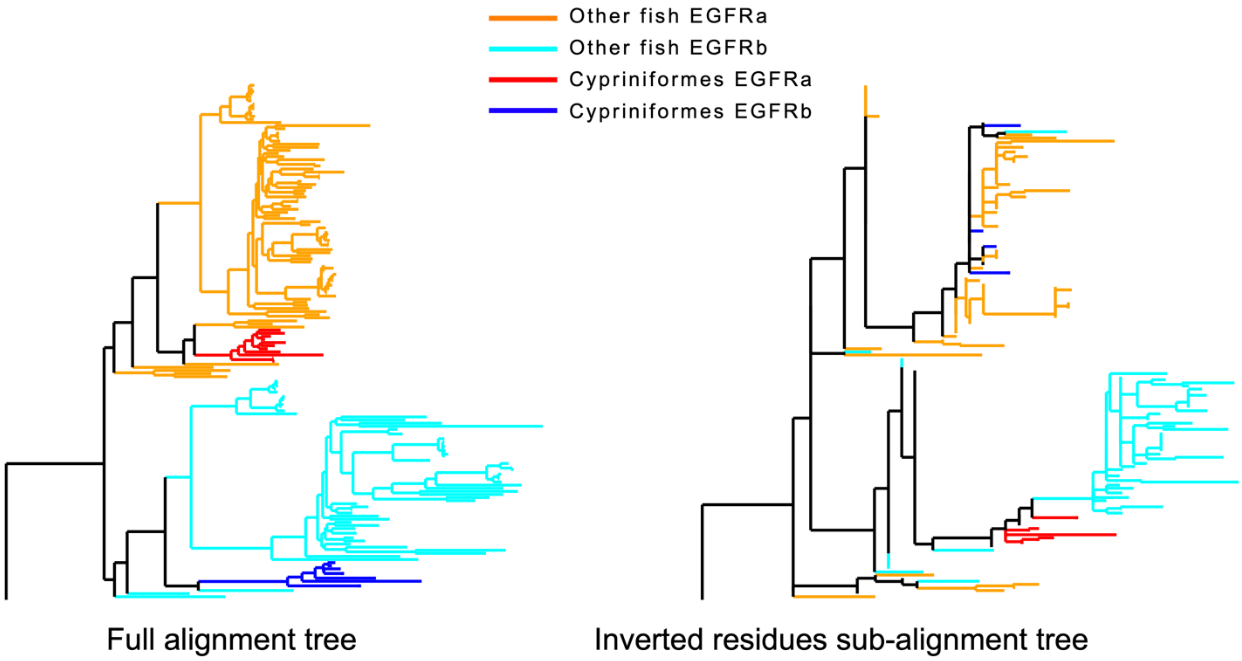

**Supplementary figure 4. Probability of a configuration.** The state of a branch (zero or one) represents whether a mutation happened in that branch. The probability of the branch state solely depends on the corresponding branch length (rate of mutation): t1 for b1 and b2, t2 for b3 to b6. The product of the six branch states gives the probability of a tree configuration. From the six branch states, it is possible to reconstruct univocally the leaf node states by counting the number of mutations in the two branches connected to a leaf node.

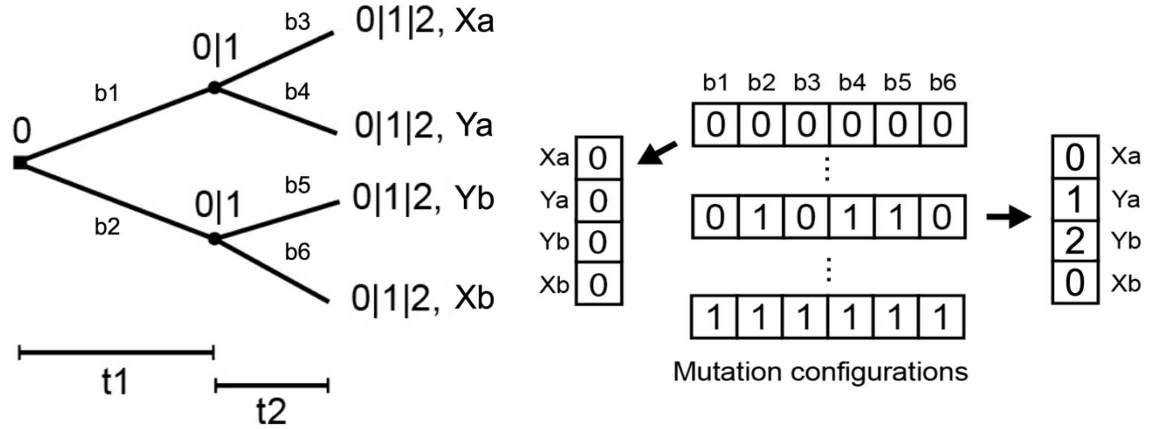

▪ Duplication node
• Speciation node

$$P_{b_i} = \begin{cases} e^{-t_x} & , if b_i = 0 \\ 1 - e^{-t_x}, & if b_i = 1 \end{cases} \qquad P_{conf} = \prod_{i=1}^{6} P_{b_i}$$

**Supplementary table 1. Fish EGFR high DIRphy scoring inverted residues.** The output of DIRphy for the sites of the phylogeny of fish EGFR in which the score is higher than the 99% percentile of the simulated evolution experiment (>0.07). The grouping of species was set to be Cypriniformes vs all other fish. "Pos" shows the residue in the reference EGFRa, "Other pos" shows the residue in the reference EGFRb. "Conservation" shows the site conservation (identity) for all EGFRa or EGFRb in the MSA jointly. The reference species for this analysis was set to *S anshuiensis*.

| pos | residue | pos_residue | other_pos | score | conservation |
|---|---|---|---|---|---|
| 104 | L | 104L | M | 0.10195107 | 0.77-0.9 |
| 137 | Q | 137Q | E | 0.25158321 | 0.74-0.73 |
| 214 | Q | 214Q | E | 0.15208071 | 0.71-0.77 |
| 253 | L | 253L | V | 0.11872061 | 0.68-0.89 |
| 275 | R | 275R | P | 0.08703647 | 0.4-0.27 |
| 292 | P | 292P | S | 0.13927292 | 0.46-0.45 |
| 309 | K | 309K | T | 0.12164597 | 0.33-0.45 |
| 387 | A | 387A | V | 0.09800167 | 0.5-0.88 |
| 423 | G | 423G | N | 0.23345058 | 0.48-0.59 |
| 438 | V | 438V | I | 0.13221518 | 0.88-0.82 |
| 491 | S | 491S | T | 0.23169011 | 0.66-0.59 |
| 506 | V | 506V | I | 0.22671569 | 0.88-0.94 |
| 520 | H | 520H | F | 0.17859498 | 0.38-0.48 |
| 558 | A | 558A | P | 0.0715249 | 0.37-0.39 |
| 609 | R | 609R | K | 0.07297936 | 0.74-0.91 |
| 811 | I | 811I | V | 0.26726795 | 0.91-0.89 |
| 967 | - | 967- | D | 0.1329542 | 0.0-0.0 |
| 1161 | - | 1161- | I | 0.136032 | 0.0-0.0 |
| 1229 | - | 1229- | T | 0.1685805 | 0.0-0.0 |
